# Differential drought sensitivity of total and active wheat rhizosphere microbiome during rainfall reduction

**DOI:** 10.64898/2026.07.08.735272

**Authors:** Abdul Samad, Ruth Lydia Schmidt, Hamed Azarbad, Julien Tremblay, Paolina Garbeva, Étienne Yergeau

## Abstract

Root-associated microorganisms play a pivotal role in helping plants adapt to drought stress. However, the underlying mechanisms of the rhizospheric microbiome under limiting soil moisture remain largely unresolved. Integrating total and active microbiome analyses enables a more accurate interpretation of microbial responses to climate change-associated water stress. We assessed the effect of reduced rainfall on two wheat genotypes, drought-tolerant (DT) and drought-sensitive (DS), using rainout shelters that allowed 100%, 75%, 50%, and 25% of natural precipitation to reach the crop. At the peak of the growing season, rhizosphere samples were collected for metagenomic (MG) and metatranscriptome (MT) sequencing. In parallel, rhizosphere volatile organic compounds (VOCs) were collected and analysed. Differential expression analysis of metatranscriptomic data using metagenomic abundance as a cofactor was performed by comparing all treatments to the 100% precipitation control. Our results demonstrate that particularly oxidative stress-related transcripts intensify in DS as rainfall decreases. Transcriptomic shifts primarily involved upregulation of transcripts associated with antioxidant (*catalase, superoxide dismutase*), heat shock proteins (*Hsp10*, *Hsp60*, *DnaK*/DnaJ, GroEL, *GroES*), as well as microbial functions related to osmoregulation, proline and glycine betaine (*PutA*, *PutP, OpuBB*), and plant growth-promoting traits such as auxin production, phosphate solubilization. Moreover, volatile organic compound (VOC) emissions differed significantly between the control and drought treatments, with higher emissions, particularly acetates, in the DS genotype than in the DT genotype. Overall, pronounced drought-induced shifts in active microbial functions and VOC emissions indicate high sensitivity and functional plasticity of the active microbiome, whereas the total microbiome remains robust under medium drought.

## Introduction

Drought driven by climate change poses a significant challenge to global food security [1]. Water deficit stress affects the plant system and its associated microbiomes at both structural and functional levels, ultimately impacting plant productivity [2]. While rhizosphere microbiomes play a pivotal role in modulating plant responses to drought, their functional dynamics under drought remain poorly understood [3]. Unlocking complex microbial responses to drought stress, particularly at the root-soil interface, is crucial for developing novel strategies to enhance drought resilience in a changing climate.

Drought-driven changes in the rhizosphere environment have been shown to influence the microbiome, often favouring drought-adapted gram-positive bacteria, such as *Actinomycetota* with strong cell walls and spore-forming ability [2, 4]. These microbes, in turn, can mediate plant stress response via synthesis of exopolysaccharides (EPS), phytohormones, volatile organic compounds, regulation of stress-responsive genes, inducing systemic tolerance, osmolytes and antioxidants production, and changing root morphology toward stress-adaptation [2, 5, 6]. Certain microbes can modulate plant stress responses by producing a variety of osmolytes, including proline, trehalose, polyamines, and amino acids [7]. Microbes can also assist plants in regulating oxidative stress by inducing the production of signalling molecules such as Jasmonic Acid (JA), Salicylic Acid (SA) [8]. These adaptations to drought can have a pronounced impact on the overall microbial activities, proliferation, and communication in the rhizosphere [9, 10]. Moreover, at the metabolomic level, water-limited conditions also influence the production and exchange of volatile metabolites at the root-soil interface. VOCs, such as terpenes, acetates and other small molecules, play essential roles in signalling between roots and microbes, modulating stress responses, and impacting microbial community functions [11, 12]. Microbial synthesis of volatile compounds can trigger plant genes involved in osmoprotectant biosynthesis (Lopes et al., 2024; Paul et al., 2022) and also alter the regulatory mechanisms of stress-responsive genes in plants [2]. These microbial mechanisms complement plant responses and together shape plant adaptation strategies under water limitation [2, 13].

Drought-induced microbial responses have been attributed to changes in both plant physiological and biochemical traits, influencing the composition of root exudates and carbon compound efflux [14, 15]. Crop varieties with different levels of drought tolerance led to differential recruitment of rhizosphere microbial taxa, with *Pseudomonadota* often dominant in the rhizospheres of drought-tolerant plants and *Actinomycetota* members preferentially associated with drought-sensitive genotypes [15, 16]. However, the mechanisms and functional implications of these changes to plants under drought stress remain poorly understood.

Our previous work on wheat holobiont demonstrated that microbial partners showed the most pronounced transcriptomic changes in response to decreasing precipitation [17]. Building on our earlier findings, this study focuses specifically on the rhizosphere microbiome to deepen understanding of its role in wheat’s adaptation to drought. An integrated multi-omics approach, including metagenomics, metatranscriptomics, and VOC profiling, was applied to the wheat rhizosphere under reduced rainfall to resolve total and active microbial genes and their associated VOCs at molecular and metabolic levels. Microbial responses under water limitation can result from shifts in community composition reflected in the metagenome, as well as from changes in gene expression within community members.

Integrating metagenomic and metatranscriptomic analyses allows these mechanisms to be better distinguished. We hypothesized that reduced precipitation would primarily impact the functional activity of the active microbiome and that these effects would vary between drought-tolerant (DT) and drought-sensitive (DS) wheat genotypes, leading to shifts in rhizosphere volatile organic compound (VOC) emissions. Wheat was chosen due to its economic and ecological significance as a staple food crop, as well as its susceptibility to drought [18]. Our approach integrated multi-omics techniques to examine the drought-induced alterations in wheat rhizosphere microbiomes under field conditions. This study lays a foundation for understanding microbial functions that strengthen rhizosphere-mediated crop resilience to climate change-associated water stress.

## Materials and Methods

### Experimental Design and Sampling

In 2016, a multi-year rainfall manipulation experiment was established at the Armand-Frappier Santé Biotechnologie Centre (Laval, Québec, Canada). The setup included rainout shelters that allowed 25%, 50%, 75%, or 100% of natural rainfall to pass through, simulating different levels of drought (Fig. 1) [19]. Each shelter covered a 2 × 2 m area and was constructed using varying numbers of transparent plastic sheets (nine, six, three, or none, respectively). The collected rainwater was directed into 20 L buckets via gutters and downspouts, and the buckets were manually emptied when full. Two wheat genotypes were planted beneath these shelters: the drought-sensitive *Triticum aestivum* cv. AC Nass and the drought-tolerant *Triticum turgidum* spp. *durum* cv. Strongfield. The design was replicated in six randomized blocks, totaling 48 plots (4 rainfall treatments × 2 genotypes × 6 blocks) (Fig. S1). Wheat was sown at a density of 500 seeds/m² with seeds harvested and replanted in the same plots the following year. Rhizosphere samples were collected in 2019 by uprooting the plant from each plot, avoiding the plot edge, and the soil loosely attached to the roots was removed. Rhizosphere soil (firmly attached soil) was collected in sterile 1.5 ml tubes. Roots were washed, separated, and stored in 15 ml sterile Falcon tubes. Samples were flash-frozen in liquid nitrogen within 2 minutes to preserve RNA integrity and stored at - 80 °C until RNA extraction.

**Figure 1.**
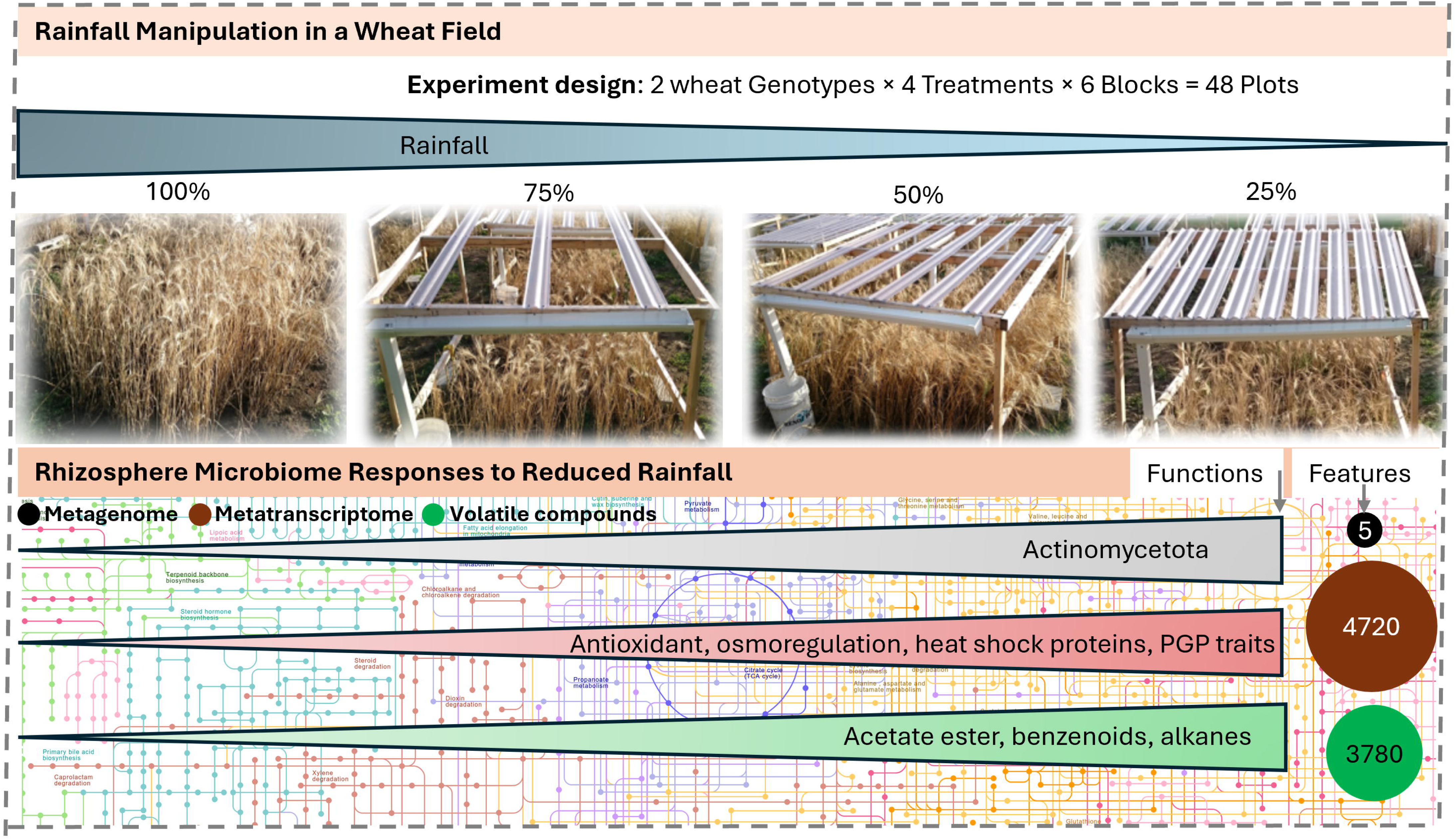
Field experiment design showing precipitation manipulation under four rainfall treatments (25%, 50%, 75%, and 100%) applied to two wheat genotypes: a drought-sensitive (DS) and a drought-tolerant (DT) genotype. Microbial responses to reduced rainfall were assessed using a multi-omics approach integrating metagenomics, metatranscriptomics, and volatile organic compound (VOC) profiling.

### Nucleic Acids Extraction and Sequencing

DNA and RNA were co-extracted from 2 g of rhizosphere soil using the RNeasy PowerSoil Total RNA Kit with the RNeasy PowerSoil DNA Elution Kit (QIAGEN, Canada). In the RNA extracts, DNA contamination was removed using DNase (Thermo Fisher Scientific, Canada), and the absence of DNA was confirmed by 16S rRNA PCR. Metagenomic and metatranscriptomic sequencing were performed on the Illumina HiSeq4000 platform (2 × 100 bp) at the Centre d’Expertise et de Services Génome Québe. Raw data are archived under NCBI Bioproject accession PRJNA1461149.

### Bioinformatics

Shotgun metagenomic libraries (2 × 150 bp) were processed in ShotgunMG [20, 21] that was developed over the GenPipes workflow management system [22]. Sequencing adapters were first removed from each read and bases at the end of reads having a quality score <30 were cut off (Trimmomatic v0.39; [23]) and scanned for sequencing adapters contaminants reads using BBDUK (BBTools v38.1) (“BBMap” n.d.) to generate quality controlled (QC) reads. The QC-passed reads from each sample were co-assembled using MEGAHIT v1.2.9 [24] with iterative kmer sizes of 31, 41, 51, 61, 71, 81, 91, 101, 111, 121 and 131 bases. Gene prediction was performed by calling genes on each assembled contig using Prodigal v2.6.3 [25]. Assignment of KEGG orthologs (KO) was done by using HMMER v3.3.2 (Johnson et al., 2010) to compare each predicted gene amino acids sequence of the co-assembly against the KOFAM database [26, 27]. The QC-passed reads were mapped (BWA mem v0.7.17; [28]) against contigs to assess quality of metagenome assembly and to obtain contig abundance profiles. Alignment files in bam format were sorted by read coordinates using samtools v1.9 [29], and only properly aligned read pairs were kept for downstream steps. Each bam file (containing properly aligned paired-reads only) was analyzed for coverage of contigs and predicted genes using bedtools (v2.31.0; [30]) using a custom bed file representing gene coordinates on each contig. Only paired reads both overlapping their contig or gene were considered for gene counts. Coverage profiles of each sample were merged to generate an abundance matrix (rows = contig, columns = samples) for which a corresponding CPM (Counts Per Million–normalized using the TMM method; edgeR v3.10.2;[31]). Taxonomic summaries were performed using a combination of in-house Perl and R scripts and Qiime v.1.9.1 [32].

Metatranscriptomic libraries (2x100) from 48 rhizosphere samples were processed for QC the same way described for metagenomic libraries processing. Quality controlled reads were then aligned against the co-assembly generated from the rhizosphere metagenomic workflow using BWA mem. Resulting alignment bam files were processed exactly as described for the metagenomic workflow to generate a metatranscriptomic gene expression matrix.

### Volatile Collection and Analysis

VOCs were collected with Rotilabo®-silicone tubes (PDMS tubes, Carl Roth GmbH & Co. KG, Karlsruhe, Germany), which were placed in each plot (see Fig. S1). The PDMS tubes were pre-treated as described previously [33]. After 20 min, PDMS tubes were removed and kept at -20◦C until analysis.

VOCs were desorbed from the PDMS tubes by using an automated thermodesorption unit (model UnityTD-100, Markes International Ltd., U.K.) at 250 ◦C for 12 min (flow 50 ml/min). The desorbed VOCs were subsequently collected on a cold trap at 10 ◦C and introduced into a GC-QTOF (model Agilent 7890B GC and the Agilent 7200AB QTOF, U.S. A.) by heating the cold trap for 10 min to 280 ◦C. A split ratio was set to 1:10. The column used was a 30 × 0.25 mm ID DB-5MS with as film thickness of 0.25 μm (Agilent 122–5532, U.S.A.). The temperature program was as follows: 2 min at 39 ◦C, 3.5 ◦C/min to 95 ◦C, 4 ◦C/min to 165 ◦C and finally 15 ◦C/min to 280 ◦C that was hold for 15 min. VOCs were detected by the MS operating at 70 eV in EI mode. Mass spectra were acquired in full scan mode (30–400 AMU, 4 spectras/s). Collected GC/MS data was converted to mzData files using the Chemstation B.06.00 (Agilent Technologies, Santa Clara, U.S.A.) and further processed (peak picking, baseline correction and peak alignment) in an untargeted manner with MZmine 2 [34]. Detected compounds were identified using NIST-MS Search by comparing the spectra, accurate mass, linear retention indices and spectra match factor with NIST 2014 V2.20 (National Institute of Standards and Technology, U.S.A., http://www.nist.gov), Wiley 9th edition, and in-house spectral libraries. Putative compounds were identified using AMDIS 2.72 (National Institute of Standards and Technology, Gaithersburg, U.S.A.) and linear retention indices of VOCs were calculated according to the method described by [35]. PDMS contaminants were removed from the final list.

### Statistical Analyses

All statistical analyses were carried out in R version 4.4.2 [36]. Differential gene abundance and differential VOC production between the treatments (75%, 50%, and 25% precipitation) and control (100% precipitation) were performed using the R package DESeq2 [37]. Differential transcript expression was assessed using the R package MTXmodel, adapted from MaAsLin2, which enables multivariable modeling of metatranscriptomic data by incorporating metagenomic abundances (gene copy numbers) as feature-specific continuous covariates in the models for differential expression analysis [38].

## Results

### Robustness of the total microbiome under rainfall reduction

A total of 48 metagenomic samples were analyzed to investigate microbial community responses to decreasing rainfall. Summary statistics for read processing, quality control, and alignment metrics are presented in Table S1. Among the wheat rhizosphere soil genes, 90.5% were bacteria, 5.6% were archaea, and 3.7% were unclassified at the kingdom level (Fig. 2A). Taxonomic analysis of metagenomes revealed stable community composition across rainfall treatments, with no major shifts in the relative abundance of dominant taxa (Fig. 2B). *Pseudomonadota* and *Acidobacteriota* were the leading phyla, followed by *Actinomycetota*. Similarly, differential abundance analysis at the gene level revealed minimal differences, with only five genes showing significant differences between the treatment and control groups (Fig. 2C, Table S2). The majority (80%) of differentially abundant genes (DAGs) were affiliated with *Actinomycetota* and encoded for hypothetical proteins.

**Figure 2.**
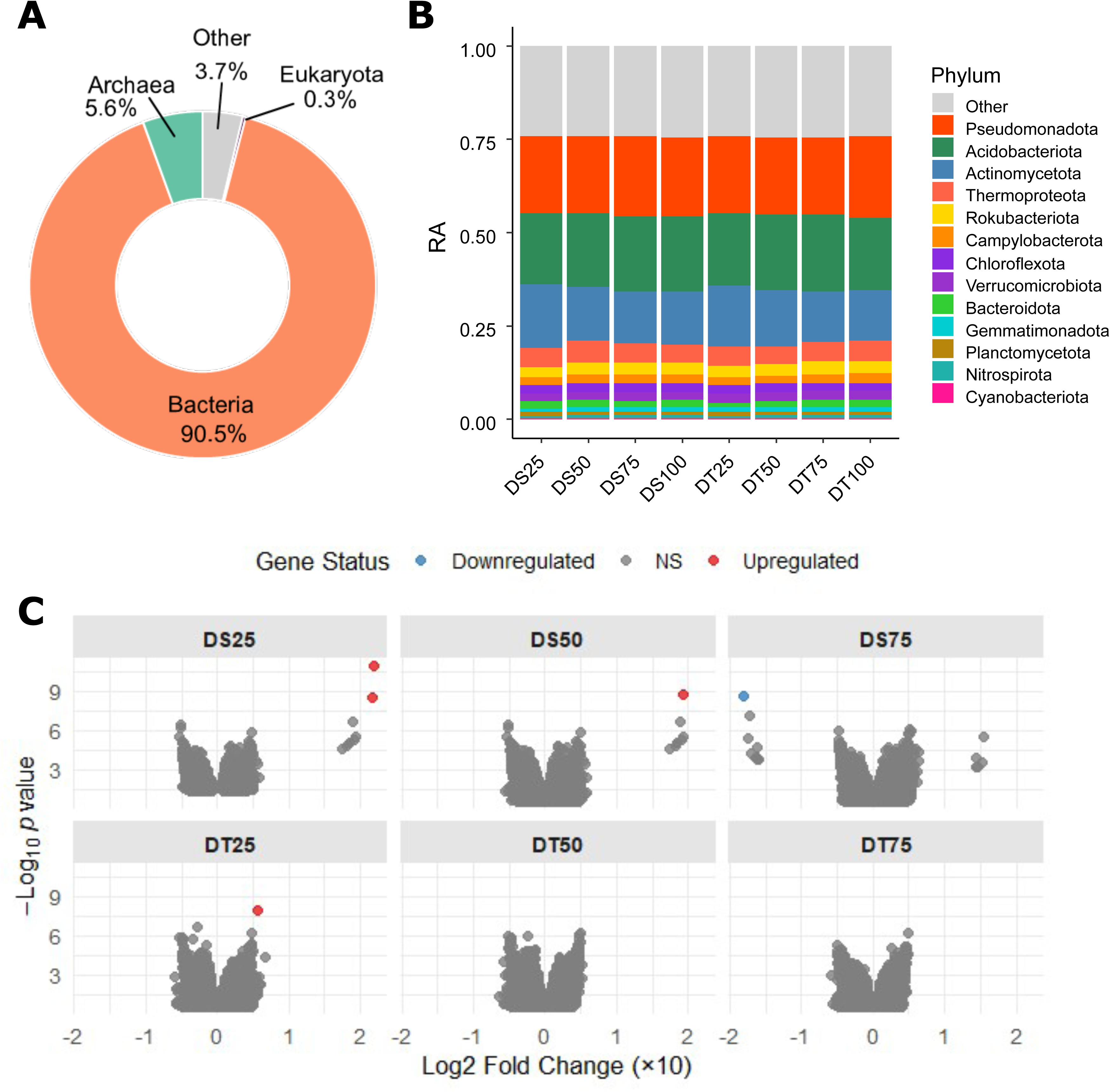
Taxonomic profile and differentially abundant genes of metagenomic data. (A) Kingdom-level relative abundance of microbial community. (B) Relative abundance (RA) of microbial community at the phylum level. (C) Differentially abundant genes between control and treatments, where all reduced rainfall treatments were compared with 100% rainfall control. NS denotes genes with no statistically significant differences.

### Dynamic shifts in the active microbiome under rainfall reduction

Differentially abundant transcripts (DATs) were identified by comparing all treatments to the 100% precipitation control. The metatranscriptome showed a stronger response to rainfall reduction than the metagenome, with an average of 788 DATs across all treatments (Fig. 3A). *Pseudomonadota*, *Acidobacteriota*, and *Actinomycetota* showed the highest number of DATs. Overall, an average of 35% of DATs were upregulated and 65% downregulated. However, more than 53% of *Actinomycetota* transcripts were upregulated under high stress in DS (DS25), whereas *Pseudomonadota* and *Acidobacteriota* exhibited a greater proportion of downregulated transcripts (Fig. 3B). Upregulated *Actinomycetota* transcripts decreased with increasing rainfall. In the DS genotype, upregulated transcripts showed increasing statistical significance (as indicated by decreasing p-values) with decreasing rainfall, reflecting more robust expression responses (Fig. 3C).

**Figure 3.**
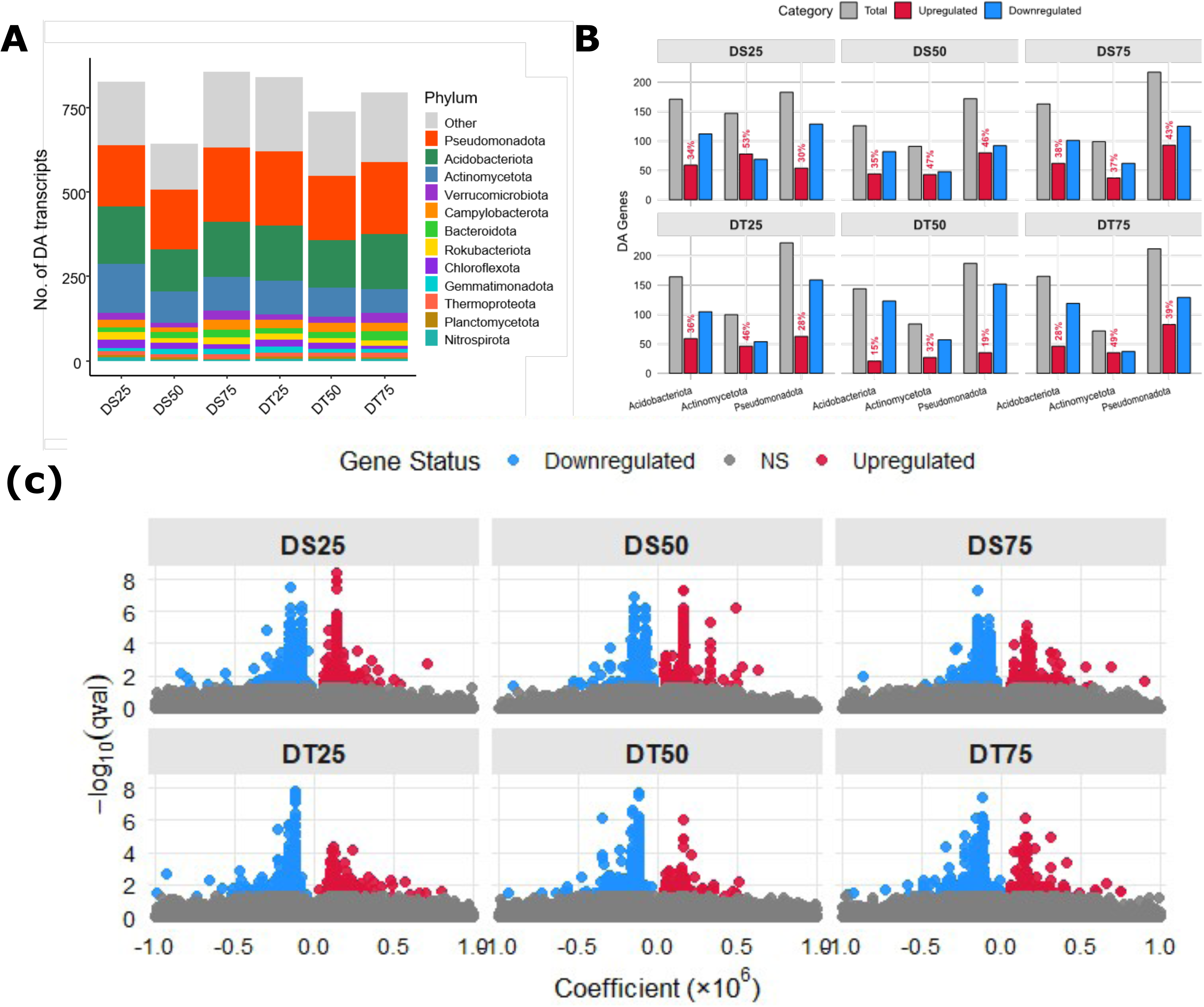
Differentially abundant transcripts (DATs) and their taxonomic origins. **(A)** Total number of DATs at each treatment with their taxonomy at the phylum level. (B) Number of upregulated and downregulated DATs for the three most abundant phyla (C). Differentially abundant transcripts between control and treatments, where all treatments were compared with 100% rainfall control. NS denotes genes with no statistically significant differences.

### Functional landscape of differentially abundant transcripts

COG analysis showed that the most differentially expressed transcripts were assigned to amino acid transport and metabolism, energy production and conversion, and carbohydrate transport and metabolism, with all three categories exhibiting a high proportion (>50%) of downregulated transcripts under drought (Fig. 4A). In contrast, the post-translational modification, protein turnover, and molecular chaperone (PTM) category displayed more than 50% upregulated transcripts and showed a clear increasing trend in transcript upregulation with decreasing precipitation. Other categories that showed elevated proportions of upregulated transcripts, particularly in the DS genotype, included inorganic ion transport and metabolism and Lipid metabolism, indicating enhanced ion homeostasis and membrane-associated adjustments under drought. Under the lowest rainfall treatments (DS25 and DT25), COG0459 (GroEL; HSP60), COG0234 (GroES; HSP10), and COG0542 (ClpA) emerged as the dominant PTM components (Fig. 4B). Notably, HSP-associated COGs (GroEL/GroES) were more dominant in DS25, indicating a stronger chaperone-mediated stress response in the rhizosphere microbial communities of this genotype. In contrast, energy production and conversion declined as drought intensified.

**Figure 4.**
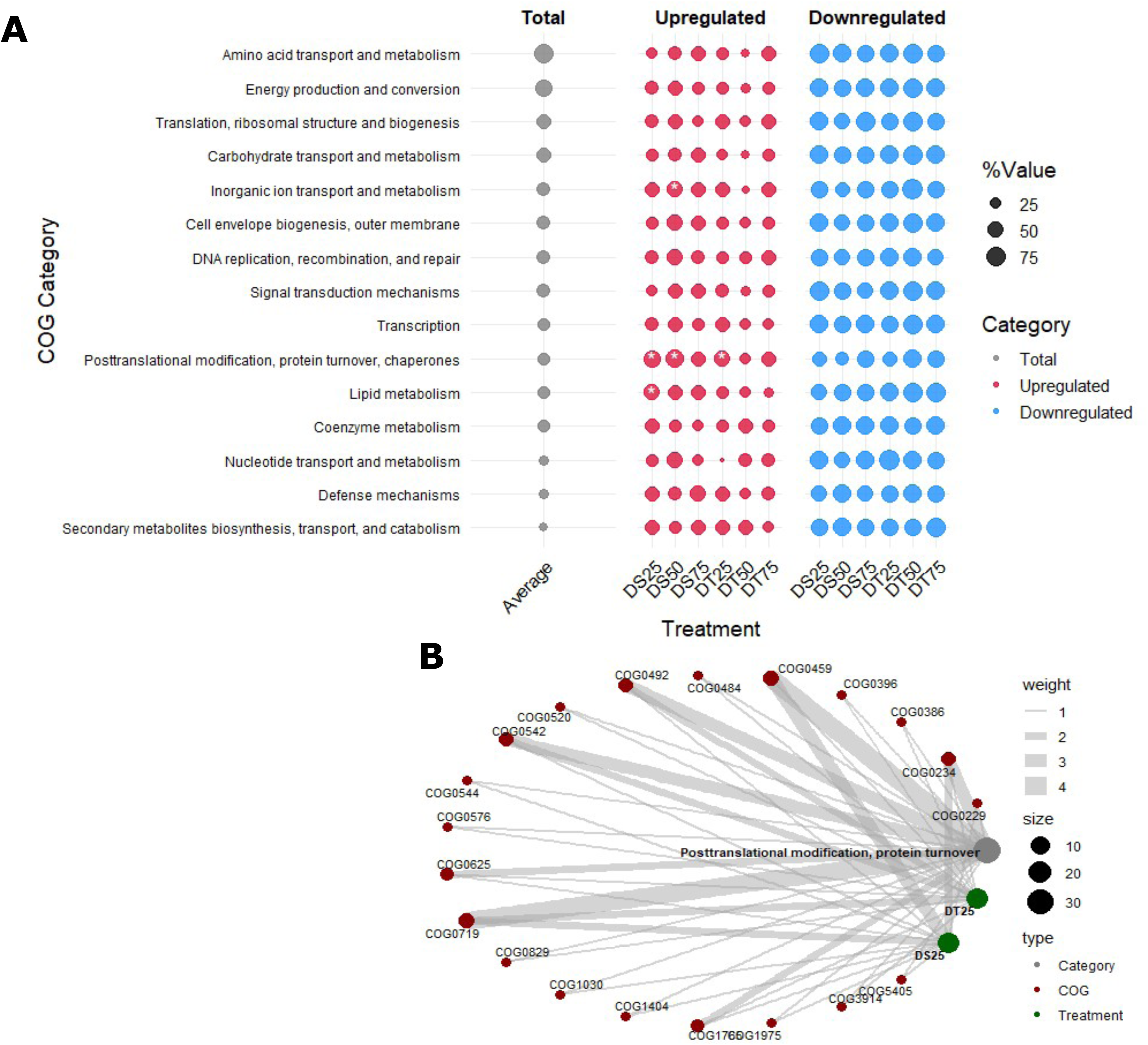
Functional annotation of differentially abundant transcripts (DATs). (A) The Clusters of Orthologous Genes (COG) categories of differentially abundant transcripts (DATs). (B) The number of COGs of upregulated transcripts assign to category post-translational modification, protein turnover, and molecular chaperone for DS and DT cultivar under 25% rainfall. The asterisk (*) denotes treatments in which more than 50% of genes were upregulated in a given COG category.

### Expression of drought-responsive and plant growth-promoting genes

Antioxidant-defense related upregulated transcripts were most abundant in DS25, followed by moderate representation in DS50 and DS75, and showed more variable patterns across the DT treatments (Fig. 5A, Table S3). Key oxidative-stress enzymes, including superoxide dismutase, catalases, glutathione S-transferases/peroxidases, cytochrome P450s, flavoprotein dehydrogenases, and several NADH: ubiquinone oxidoreductase subunits were especially enriched in DS25. Molecular chaperone and post-translational modification (PTM) functions were enriched under 25% rainfall treatments. ATP-binding subunits of the Clp protease, DnaK/DnaJ chaperones, and the chaperonin GroEL (HSP60) were consistently upregulated in both DS25 and DT25, with a stronger response observed in DS25 (Fig. 5B). Osmoregulation-related functions showed distinct patterns, exhibiting the upregulated transcripts, including proline and glycine betaine transport system, choline dehydrogenase, glucose dehydrogenase, glucose/sorbosone dehydrogenases, and sugar-processing enzymes such as alpha-L-arabinofuranosidase and dTDP-D-glucose 4,6-dehydratase. Transport-related systems, reflecting microbial adaptation to mild osmotic stress at 75% rainfall (Fig. 5C). Moreover, plant growth-promoting (PGP) traits were also upregulated in the rhizosphere under drought stress. Quorum-sensing-related upregulated transcripts were found across all treatments, while auxin (IAA) production was detected exclusively in DS25, and phosphate solubilization genes (*alkaline phosphatase D*) were observed in DS50 and DT25 (Fig. 5D).

**Figure 5.**
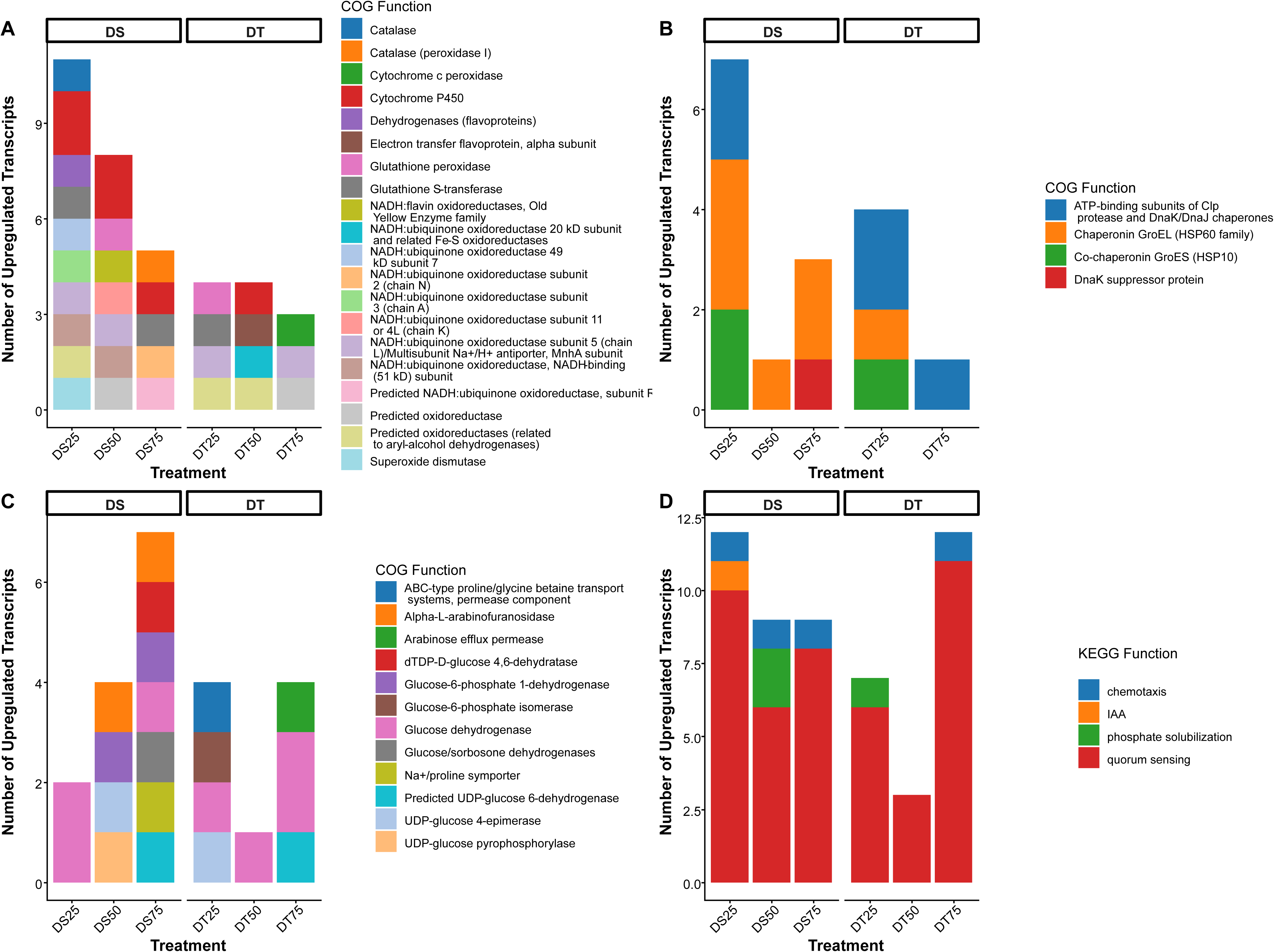
Upregulated differentially abundant transcripts (DATs) linked to drought stress and plant performance. (A) upregulated transcripts assigned to antioxidant defense, (B) upregulated transcripts assigned to heat shock proteins, (C) upregulated transcripts assigned to osmotic adjustment, (D) upregulated transcripts related to plant performance.

### Transcriptomic functions shared across genotypes and treatments

Across all genotypes and drought treatments, five COGs of upregulated transcripts were consistently shared, including DNA replication and repair, signal transduction and motility, enzymatic processing of metabolites, and intracellular signalling. Specifically, these included COG0577, involved in antimicrobial peptide transport; COG0587, encoding DNA polymerase III; COG2204, a motility- and chemotaxis-related response regulator with CheY-like receiver, AAA-type ATPase, and DNA-binding domains; COG0596, predicted hydrolases or acyltransferases of the alpha/beta hydrolase superfamily; and COG2114, an adenylate cyclase producing intracellular signalling molecules (Fig. 6A, Table 1). Together, these conserved functions likely support microbial adaptation and resilience under drought stress. Across DS treatments, 24 COGs were consistently shared, reflecting core microbial functions under drought stress. Among these, COG2197 (response regulator containing a CheY-like receiver domain and an HTH DNA-binding domain), COG2124 (cytochrome P450), and COG0515 (Serine/threonine protein kinase) were particularly noteworthy (Fig. 6B, Table 1). In DT treatments, 15 COGs were uniquely shared, reflecting drought-responsive microbial functions specific to this genotype. In particular, COG4993 (glucose dehydrogenase) suggests enhanced energy metabolism and redox balance, COG1680 (beta-lactamase class C and penicillin-binding proteins) indicates cell wall remodelling and structural adaptation under stress, and COG0642 (signal transduction histidine kinase) highlights environmental sensing and regulatory responses (Fig. 6C and Table 1).

**Figure 6.**
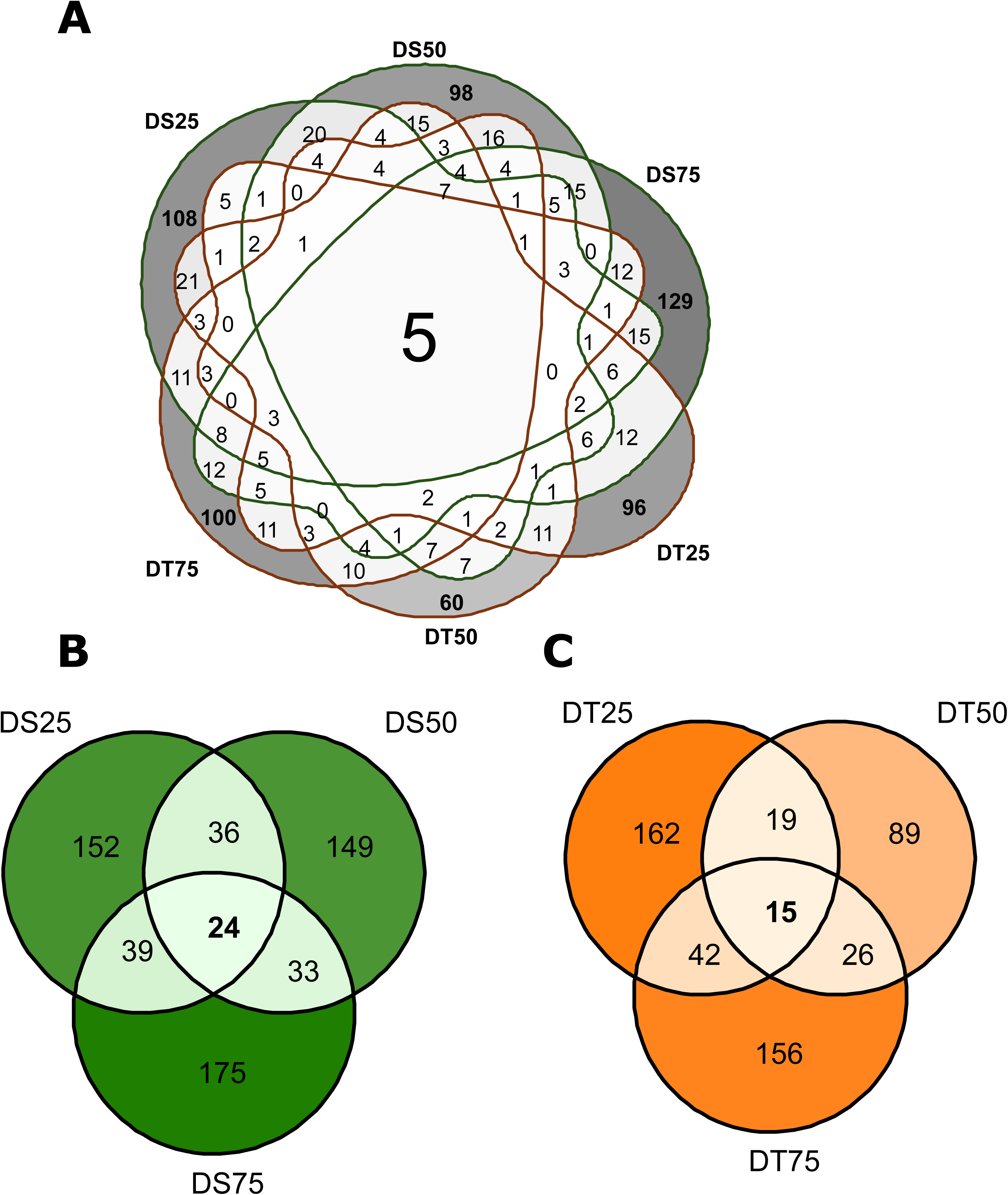
Shared and group-specific COG assigned to upregulated transcripts. **(A)** Number of COGs shared between treatments and cultivars, (B) Number of COGs shared between DS treatments, (C) Number of COGs shared between DT treatments. The functions of shared COGs are shown in Table 1.

**Table 1.**
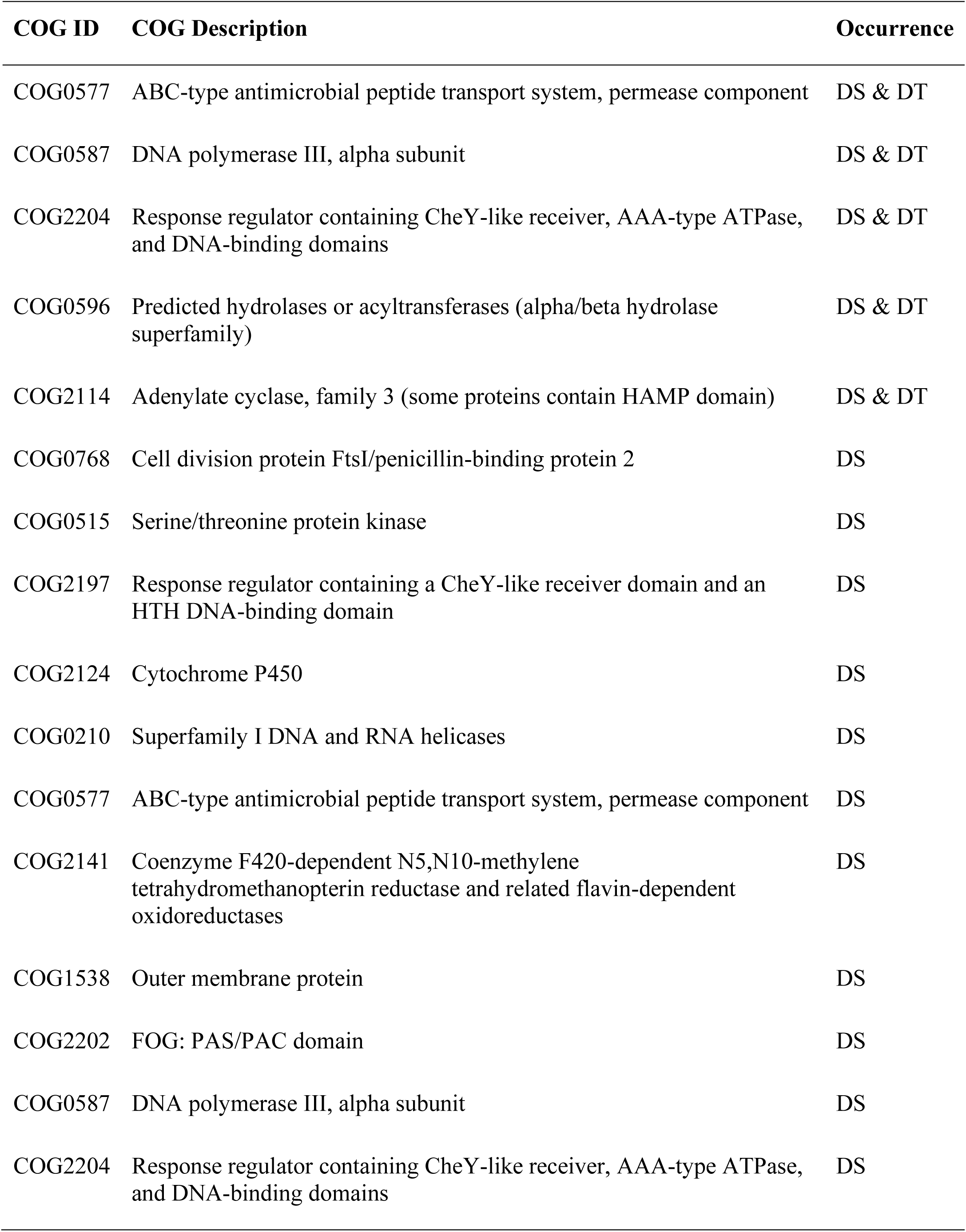
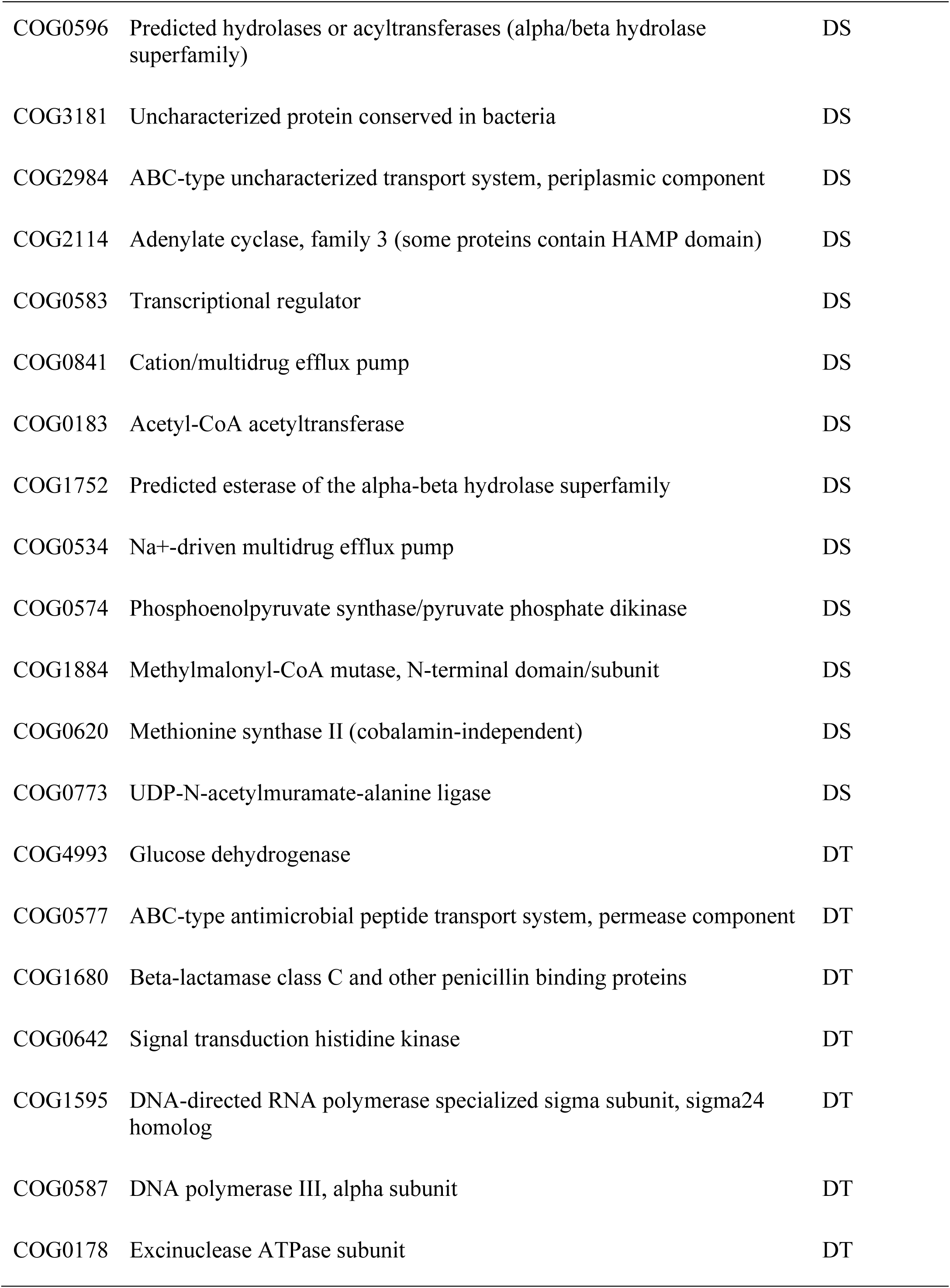
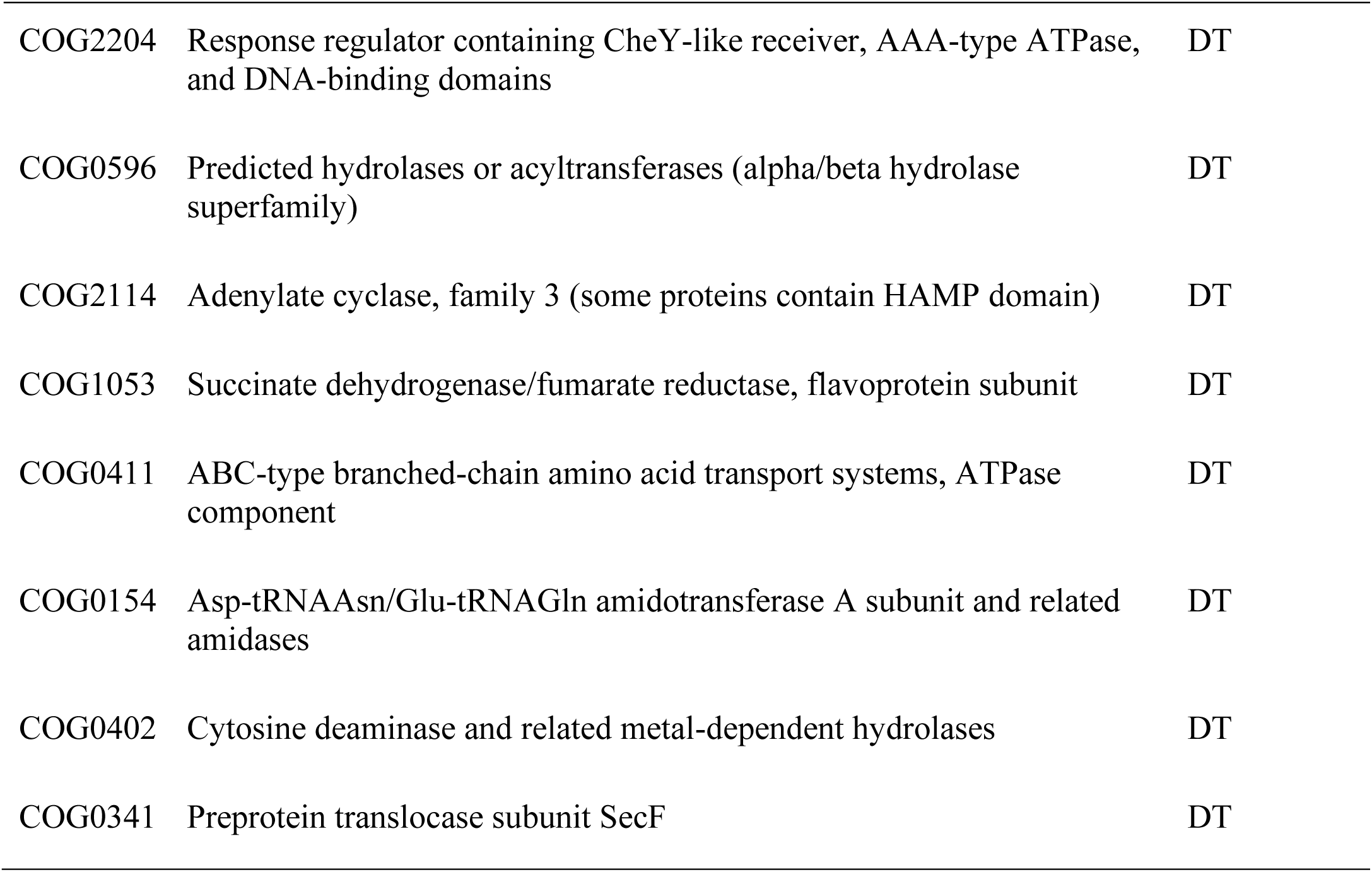
Shared COG functional categories across treatments and genotypes.

| COG ID | COG Description | Occurrence |
| --- | --- | --- |
| COG0577 | ABC-type antimicrobial peptide transport system, permease component | DS & DT |
| COG0587 | DNA polymerase III, alpha subunit | DS & DT |
| COG2204 | Response regulator containing CheY-like receiver, AAA-type ATPase, and DNA-binding domains | DS & DT |
| COG0596 | Predicted hydrolases or acyltransferases (alpha/beta hydrolase superfamily) | DS & DT |
| COG2114 | Adenylate cyclase, family 3 (some proteins contain HAMP domain) | DS & DT |
| COG0768 | Cell division protein FtsI/penicillin-binding protein 2 | DS |
| COG0515 | Serine/threonine protein kinase | DS |
| COG2197 | Response regulator containing a CheY-like receiver domain and an HTH DNA-binding domain | DS |
| COG2124 | Cytochrome P450 | DS |
| COG0210 | Superfamily I DNA and RNA helicases | DS |
| COG0577 | ABC-type antimicrobial peptide transport system, permease component | DS |
| COG2141 | Coenzyme F420-dependent N5,N10-methylene tetrahydromethanopterin reductase and related flavin-dependent oxidoreductases | DS |
| COG1538 | Outer membrane protein | DS |
| COG2202 | FOG: PAS/PAC domain | DS |
| COG0587 | DNA polymerase III, alpha subunit | DS |
| COG2204 | Response regulator containing CheY-like receiver, AAA-type ATPase, and DNA-binding domains | DS |
| COG0596 | Predicted hydrolases or acyltransferases (alpha/beta hydrolase superfamily) | DS |
| COG3181 | Uncharacterized protein conserved in bacteria | DS |
| COG2984 | ABC-type uncharacterized transport system, periplasmic component | DS |
| COG2114 | Adenylate cyclase, family 3 (some proteins contain HAMP domain) | DS |
| COG0583 | Transcriptional regulator | DS |
| COG0841 | Cation/multidrug efflux pump | DS |
| COG0183 | Acetyl-CoA acetyltransferase | DS |
| COG1752 | Predicted esterase of the alpha-beta hydrolase superfamily | DS |
| COG0534 | Na<sup>+</sup>-driven multidrug efflux pump | DS |
| COG0574 | Phosphoenolpyruvate synthase/pyruvate phosphate dikinase | DS |
| COG1884 | Methylmalonyl-CoA mutase, N-terminal domain/subunit | DS |
| COG0620 | Methionine synthase II (cobalamin-independent) | DS |
| COG0773 | UDP-N-acetylmuramate-alanine ligase | DS |
| COG4993 | Glucose dehydrogenase | DT |
| COG0577 | ABC-type antimicrobial peptide transport system, permease component | DT |
| COG1680 | Beta-lactamase class C and other penicillin binding proteins | DT |
| COG0642 | Signal transduction histidine kinase | DT |
| COG1595 | DNA-directed RNA polymerase specialized sigma subunit, sigma24 homolog | DT |
| COG0587 | DNA polymerase III, alpha subunit | DT |
| COG0178 | Excinuclease ATPase subunit | DT |
| COG2204 | Response regulator containing CheY-like receiver, AAA-type ATPase, and DNA-binding domains | DT |
| COG0596 | Predicted hydrolases or acyltransferases (alpha/beta hydrolase superfamily) | DT |
| COG2114 | Adenylate cyclase, family 3 (some proteins contain HAMP domain) | DT |
| COG1053 | Succinate dehydrogenase/fumarate reductase, flavoprotein subunit | DT |
| COG0411 | ABC-type branched-chain amino acid transport systems, ATPase component | DT |
| COG0154 | Asp-tRNA<sup>Asn</sup>/Glu-tRNA<sup>Gln</sup> amidotransferase A subunit and related amidases | DT |
| COG0402 | Cytosine deaminase and related metal-dependent hydrolases | DT |
| COG0341 | Preprotein translocase subunit SecF | DT |

### Differential production of VOCs under rainfall reduction

Across treatments and controls, a large number of volatile organic compounds (VOCs) were differentially abundant, reflecting dynamic microbial and plant responses to drought (Table S4, Fig. 7A). Volatiles with increased emission were most pronounced in DS25, where they accounted for the highest percentage of total differentially abundant VOCs, and gradually declined in DS50 and DS75 (Fig. 7B). In contrast, DT treatments showed fewer differentially abundant VOCs compared to DS treatments. The dominant rhizosphere VOCs identified were toluene, 1-butanol, 3-methyl-, acetate, pentane, 3-methyl-, 1-tridecyne, and Trp-Met (Table 2, Fig. 7A). Toluene exhibited consistent emissions in both DS and DT treatments, suggesting stable and constitutive production irrespective of drought intensity. In contrast, 1-butanol, 3-methyl-, acetate and 1-tridecyne were abundant in DS treatments, while pentane, 3-methyl- was predominantly associated with DT treatment. Overall, the detected VOCs comprised diverse chemical classes, including esters, organic acids, and aromatic compounds, with DS and DT treatments exhibiting distinct responses under reduced rainfall conditions (Table S5).

**Figure 7.**
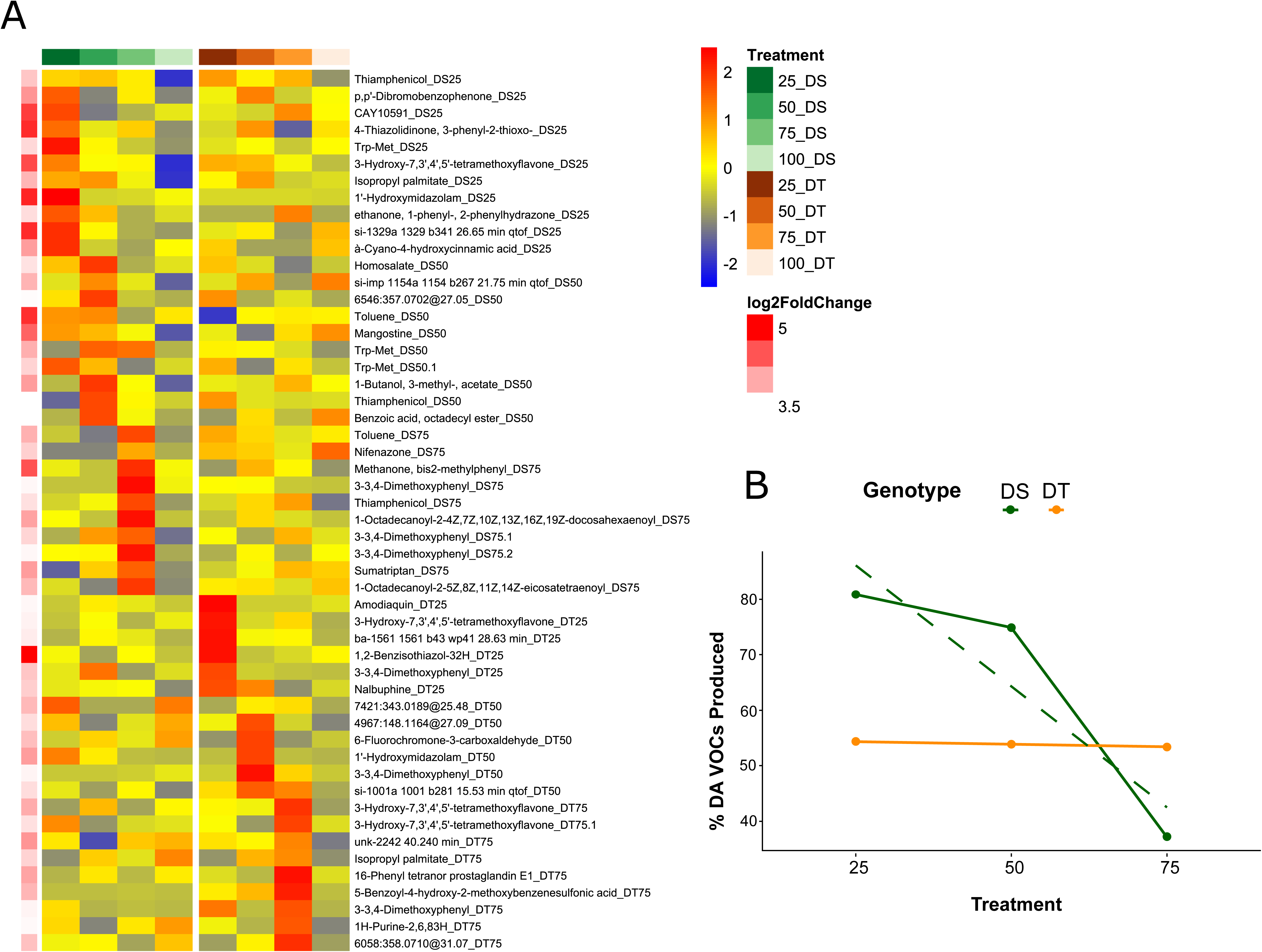
Volatile organic compounds (VOCs). (A) Heatmap of significantly produced volatile organic compounds (VOCs) in drought-sensitive (DS) and drought-tolerant (DT) wheat genotypes under decreasing rainfall treatments. Only VOCs with log2 fold change > 3 and p < 0.05 relative to the 100% rainfall control were included. Rows represent VOCs and columns represent mean normalized treatment abundances. Colors indicate row-wise scaled abundance (red, higher; blue, lower). Right-side annotations show log2 fold change values. VOC labels ending with treatment identifiers (e.g., Thiamphenicol_DS25) indicate the treatment in which the VOC was significantly enriched (B) percentage of VOCs with significantly increased production among all differentially abundant VOCs, (C).

**Table 2.** Detected Compounds previously reported as microbial volatile organic compounds in the mVOC 4.0 database.

| Compound | Treatment | RT | log2FoldChange | pvalue |
| --- | --- | --- | --- | --- |
| 1-Butanol, 3-methyl-, acetate | DS25 | 2.77 | 0.662 | 0.032 |
| 1-Butanol, 3-methyl-, acetate | DS25 | 2.76 | 1.436 | 0.034 |
| 1-Butanol, 3-methyl-, acetate | DS50 | 2.77 | 1.333 | 0.000 |
| 1-Butanol, 3-methyl-, acetate | DS50 | 2.76 | 3.550 | 0.001 |
| 1-Butanol, 3-methyl-, acetate | DS50 | 2.76 | 1.619 | 0.001 |
| 1-Butanol, 3-methyl-, acetate | DS50 | 2.76 | 1.395 | 0.015 |
| 1-Butanol, 3-methyl-, acetate | DS50 | 2.76 | 1.164 | 0.025 |
| 1-Butanol, 3-methyl-, acetate | DS75 | 2.84 | 1.198 | 0.014 |
| 1-Butanol, 3-methyl-, acetate | DS75 | 2.76 | 1.785 | 0.026 |
| 1-Butanol, 3-methyl-, acetate | DS75 | 2.76 | 1.375 | 0.043 |
| 1-Butanol, 3-methyl-, acetate | DS75 | 2.76 | 1.044 | 0.045 |
| 1-Butanol, 3-methyl-, acetate | DT25 | 2.76 | 2.078 | 0.010 |
| 1-Butanol, 3-methyl-, acetate | DT25 | 2.76 | 1.372 | 0.011 |
| 1-Butanol, 3-methyl-, acetate | DT75 | 2.76 | 1.399 | 0.015 |
| 1-Butanol, 3-methyl-, acetate | DT75 | 2.76 | 1.071 | 0.048 |
| 1-Tridecyne | DS25 | 9.69 | 0.988 | 0.011 |
| 1-Tridecyne | DS50 | 9.69 | 0.878 | 0.024 |
| 1-Tridecyne | DS75 | 9.69 | 1.094 | 0.026 |
| 1-Tridecyne | DS75 | 9.69 | 0.771 | 0.047 |
| Cyclotrisiloxane, hexamethyl- | DS25 | 4.91 | 1.006 | 0.011 |
| Cyclotrisiloxane, hexamethyl- | DS75 | 4.91 | 0.894 | 0.005 |
| Cyclotrisiloxane, hexamethyl- | DS75 | 4.89 | 1.634 | 0.018 |
| Cyclotrisiloxane, hexamethyl- | DS75 | 4.91 | 0.797 | 0.024 |
| Isobutane | DS50 | 2.27 | 0.973 | 0.008 |
| Isobutane | DT75 | 2.27 | 1.007 | 0.006 |

Wheat responses to drought stresses
| Compound | Treatment | RT | log2FoldChange | pvalue |
| --- | --- | --- | --- | --- |
| Pentane, 3-methyl- | DS75 | 2.65 | 2.169 | 0.012 |
| Pentane, 3-methyl- | DS75 | 2.65 | 1.487 | 0.044 |
| Pentane, 3-methyl- | DT25 | 2.65 | 0.942 | 0.024 |
| Pentane, 3-methyl- | DT75 | 2.65 | 1.248 | 0.003 |
| Pentane, 3-methyl- | DT75 | 2.65 | 0.749 | 0.005 |
| Pentane, 3-methyl- | DT75 | 2.65 | 1.675 | 0.023 |
| Toluene | DS25 | 4.35 | 1.976 | 0.016 |
| Toluene | DS50 | 4.35 | 3.727 | 0.001 |
| Toluene | DS50 | 4.34 | 1.193 | 0.021 |
| Toluene | DS75 | 4.33 | 4.815 | 0.000 |
| Toluene | DS75 | 4.35 | 1.906 | 0.000 |
| Toluene | DS75 | 4.34 | 1.441 | 0.000 |
| Toluene | DS75 | 4.35 | 1.474 | 0.003 |
| Toluene | DS75 | 4.36 | 1.094 | 0.003 |
| Toluene | DS75 | 4.34 | 1.105 | 0.008 |
| Toluene | DS75 | 4.35 | 1.040 | 0.014 |
| Toluene | DS75 | 4.35 | 1.362 | 0.015 |
| Toluene | DT25 | 4.35 | 2.080 | 0.000 |
| Toluene | DT25 | 4.35 | 0.607 | 0.034 |
| Toluene | DT25 | 4.35 | 0.776 | 0.045 |
| Toluene | DT25 | 4.34 | 0.679 | 0.046 |
| Toluene | DT50 | 4.35 | 1.034 | 0.008 |
| Toluene | DT50 | 4.34 | 0.597 | 0.010 |
| Toluene | DT50 | 4.35 | 1.260 | 0.024 |
| Toluene | DT50 | 4.35 | 0.632 | 0.027 |
| Toluene | DT50 | 4.35 | 0.872 | 0.040 |
| Toluene | DT75 | 4.35 | 0.787 | 0.006 |
| Toluene | DT75 | 4.35 | 0.934 | 0.012 |
| Toluene | DT75 | 4.33 | 0.791 | 0.026 |
| Toluene | DT75 | 4.35 | 1.033 | 0.039 |

## Discussion

Our study offers deeper insight into the wheat rhizosphere microbiome across varying rainfall gradients, building on our previous research that highlighted the microbiome as the most responsive component of the wheat holobiont [17]. To explore the mechanisms underlying these responses, we integrated both the total and active rhizosphere microbiome and profiled total rhizosphere VOC emissions. We hypothesized that reduced rainfall would primarily impact the functional activity of the active microbiome, with effects differing between DS and DT genotypes, potentially driving shifts in VOC production. This multi-omics approach allowed us to dissect how rhizosphere microbial activity, in combination with broader chemical signals from the rhizosphere, contributes to plant-microbe interactions and modulates stress responses under water limitation.

Pronounced transcriptional differences among treatments relative to the control suggest that rainfall reduction exerts a strong influence on microbial activity at the expression level. Transcriptional reprogramming of microbiome in response to drought is evident from the previous studies [2, 11, 17]. The upregulation of genes involved in post-translational modification, molecular chaperones, and coenzyme metabolism highlights the central role of protein stabilization, repair, and redox homeostasis in bacterial adaptation to drought (J. Li et al., 2025; Sharma et al., 2023). The enhanced expression of antioxidant defense genes (e.g., KatE) and molecular chaperones (DnaK, SufA) reflects a coordinated bacterial strategy to mitigate oxidative damage and maintain cellular integrity under water-deficit conditions (Kour & Yadav, 2022; J. Li et al., 2025). The upregulated transcript indicates elevated detoxification, with broader oxidoreductase activity helping maintain redox homeostasis under water stress. Several osmotic adjustment-related functions were consistently detected among upregulated transcripts, highlighting the microbial strategies that support rhizosphere resilience under water deficit. The activation of carbohydrate dehydrogenases and sugar-processing enzymes suggests metabolic reprogramming to support osmoprotection and redox balance under reduced soil water availability. Concurrent upregulation of compatible solute transport systems, including proline, indicates enhanced osmolyte acquisition as a microbial drought-adaptation strategy under moderate reductions in soil water content. Such osmotic adjustment is well documented in soil and rhizosphere microbiomes exposed to water limitation and reflects genotype-dependent modulation of microbial stress responses [41, 42]. The upregulated transcript of microbial auxin production, phosphate solubilization, chemotaxis and quorum sensing indicate active plant-microbe cooperation to sustain root growth and nutrient acquisition coordination to enhance rhizosphere stability and stress resilience. Such functions are widely recognized as key microbial strategies for improving holobiont fitness under fluctuating moisture conditions [43, 44]. The downregulation of energy production pathways suggests a shift towards energy conservation, a hallmark of microbial survival under environmental stress, as evidenced by long-term soil warming studies in which microbes reduced ribosomal content and translation to conserve energy [45]. Overall, the preferential upregulation of antioxidant-related functions under more severe moisture limitation reflects microbial investment in oxidative stress mitigation, whereas the dominance of osmoregulatory processes under low to moderate stress indicates adaptive adjustment to moderate water availability. Such stress-intensity dependent functional shifts are widely reported in drought-affected soil and rhizosphere microbiomes [42, 43].

The contrasting transcriptomic responses between DS and DT genotypes highlight the host’s influence on rhizosphere microbiome activation under stress [46]. The predominance of upregulated *Actinomycetota* transcripts in DS and *Pseudomonadota* transcripts in DT suggests a genotype-specific effect, consistent with prior findings that host genotype can tailor the functional adaptation of associated microbial communities under drought [16, 47]. Functionally, DS microbiomes showed mainly general stress responses, including upregulation of molecular chaperones and protein kinases, suggesting a reactive strategy to offset host vulnerability. In contrast, DT microbiomes were enriched in functions related to energy metabolism, structural adaptation, and environmental sensing, indicative of a more proactive microbial support under water limitation. These patterns suggest that differences in drought sensitivity between genotypes are partly shaped by how their rhizosphere communities respond to stress, highlighting the key role of host-microbiome interactions in plant drought tolerance [48].

However, the metagenomic (MAG) abundance showed only a weaker response across the rainfall intensities. Limited differential abundance at the DNA level suggests that moderate drought induces microbial responses primarily through RNA-level adjustments, with minimal compositional restructuring [17, 49, 50]. A similar pattern has been reported under drought conditions, where metatranscriptomic analyses showed pronounced shifts in active microbial functional and taxonomic profiles, while overall functional potential and total taxonomic diversity remained largely stable [11, 49].

Consistent with the strong metatranscriptomic response, rhizosphere volatile emissions were markedly enhanced under drought, with acetate esters and aromatic compounds emerging as dominant drought-responsive signals, particularly in DS treatments. Previous studies have shown that under drought, microbial responses to drought shift carbon cycling from biomass production and carbon storage toward the release of specific volatile metabolites, including acetate [11, 51]. Moreover, carbon cycling is likely to be impacted by increased air-filled pore space under drought, which restricts aqueous connectivity and isolates microbial communities in microhabitats, limiting access to non-volatile carbon substrates [11, 42]. In parallel, soil drying disrupts hydrological connectivity by thinning water films and draining macropores, increasing diffusion distances and limiting substrate transport to microbial cells. Reduced diffusivity imposes substrate limitation independent of direct physiological stress, whereas gas-phase VOCs can bypass these aqueous constraints, allowing metabolite exchange despite limited water-mediated transport [42]. In line with VOC production, acetate-associated metabolic functions were upregulated, with acetyl-CoA C-acetyltransferase (K00626) as the dominant enzyme. As key components of central carbon metabolism, their coordinated induction suggests a shift in carbon flux toward acetate metabolism under drought stress, potentially contributing to increased acetate-derived VOC emissions. Together, these findings indicate that drought stress drives the emission of specific volatile compounds through both metabolic reprogramming and physical constraints on substrate availability. Volatiles captured in the rhizosphere may originate from both plant roots and associated microorganisms, and distinguishing their sources is challenging because many compounds can be produced by both. However, a substantial proportion of the volatiles identified in this study are reported in the mVOC database as being of microbial origin. This highlights the combined influence of microbial metabolism, plant physiology and soil physics in shaping rhizosphere volatile profiles under water limitation. Changes in VOC emissions may contribute to plant–microbe signaling, potentially acting as a ‘cry for help’ mechanism to recruit beneficial microorganisms under stress, with potential implications for plant-microbe interactions and ecosystem carbon flux.

In conclusion, drought strongly alters the functional activity of the wheat rhizosphere microbiome, with pronounced transcriptomic and VOC responses but limited changes in community composition. These findings indicate that microbial adaptation to water limitation is primarily driven by transcriptional regulation rather than structural shifts. Identifying key drought-responsive microbial functions provides a basis to engineer drought-resilient crop microbiomes.

## Author Contributions

Abdul Samad (Performed all data analyses and wrote the manuscript), Ruth Schmidt (Collected data and contributed to VOC data analysis), Hamed Azarbad (Contributed to the experiments and reviewed the manuscript), Julien Tremblay (Conducted bioinformatics analyses, curated data, and reviewed the manuscript), Paolina Garbeva (Provided valuable input on VOC data analysis and contributed to manuscript review and editing), Etienne Yergeau (Designed the experiments, supervised the project, and contributed to manuscript review and editing).

## Data availability

Raw metagenomic and metatranscriptomic sequencing data generated in this study are available at https://dataview.ncbi.nlm.nih.gov/object/PRJNA1461149?reviewer=5u0ukltd0em3kqtvn9uq7k8r6r. Processed datasets supporting the findings of this study, together with the code used for data analysis and figure generation, are available at Zenodo https://zenodo.org/records/20775800 and GitHub https://github.com/samad2386/wheat_rhizo_microbiome.

## Figures legends

**Figure S1.**
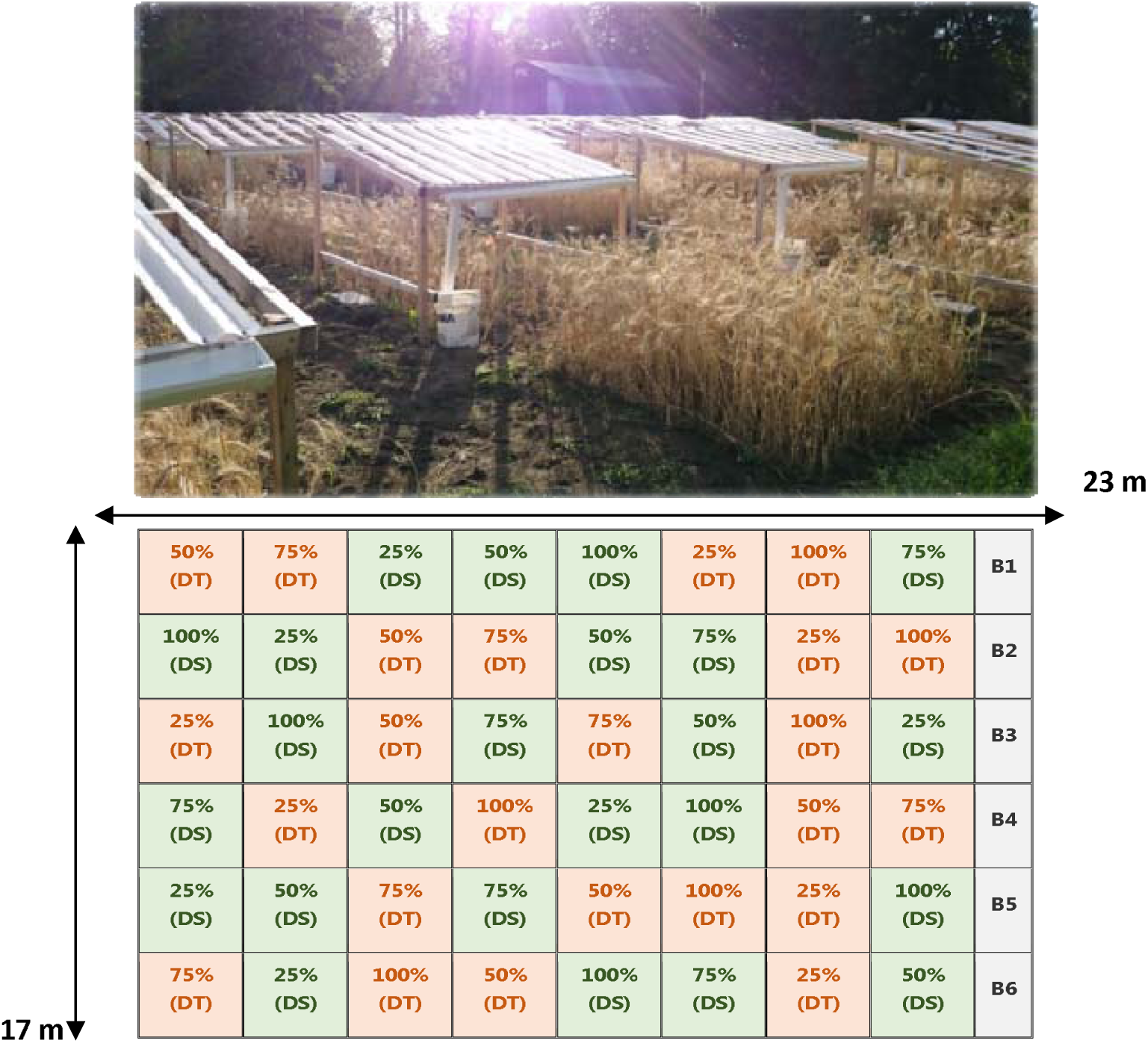
Field experiment manipulating precipitation using four rainfall treatments ( 25%, 50%, 75% and 100%) and two wheat cultivars, a drought-sensitive (DS) and a drought-tolerant (DT) cultivar, arranged in six blocks. The experiment was established in 2016 at the Armand-Frappier Santé Biotechnologie Research Center (Québec, Canada). In total, 48 plots (2 × 2 m) were arranged in a randomized complete block design. The figure was adapted from Wang et al. (2022).

